# Geographic redistributions are insufficient to mitigate the erosion of species’ environmental niches

**DOI:** 10.1101/2024.06.04.596070

**Authors:** Jeremy Cohen, Walter Jetz

## Abstract

As climate change accelerates, species may survive in place thanks to niche plasticity or adaptation or must redistribute to conserve their environmental niches^1^. Examples of such geographical range shifts abound^2–4^, but to date an assessment of species’ success in retaining niches and limiting their climate change exposure is missing. Here, we develop a novel method to account for biases inherent in tens of millions of citizen science observations, allowing us to evaluate how species have mitigated their climatic niche loss using geographical redistributions. We find that over 20 years, 384 North American bird species shifted their summertime distributions 0.73° north, mitigating their expected exposure to warming by ∼1.16 °C and thus averting 44% of expected niche loss had they stayed in place. Despite these movements, species were still exposed to an average ∼1.47 °C increase in temperature and few species achieved complete niche retention. Meanwhile, species only mitigated ∼0.47 °C (11%) in winter, shifting their niches by ∼3.74 °C, with almost no species fully retaining their niches. Species moving the furthest north succeeded most in conserving niches across both seasons. As expected, but previously untested at this scale, species that have physiological characteristics associated with dispersal achieved the greatest redistributions and niche retention. Most geographical redistributions have only been partially effective towards mitigating climate change and the gap between climate change exposure and species’ historical niches is growing rapidly even in a highly mobile group such as birds, raising concerns about the ability of less mobile taxa to persist in a warmer world.

## Main Text

Species require a specific set of environmental conditions – environmental niches – for persistence^5–7^, but are increasingly exposed to novel conditions outside of these windows as climate change intensifies^8–11^. Next to plastic and evolutionary responses^12–14^, geographical redistributions are considered the central response allowing species to limit climate change risk^15^. For species to mitigate the erosion of their climatic niches and avoid novel conditions such as warmer temperatures, redistributions towards the poles or upslope have been identified as a key mechanism for survival^1,4,16^. Such geographical range shifts have been extensively documented^2,17,18^, but the extent to which these indeed allow species to retain their historic climatic niches remains unclear^19,20^.

As climate change accelerates, connecting the links between climate change exposure, geographical redistributions, and climatic niche retention is critical^4,15,20,21^. Many projections assume that species will fully track their climatic niches over time, even over thousands of kilometers^22^, though this is likely unrealistic given dispersal limitations^23^ and the rapid pace of ongoing climate change^24^. However, smaller redistributions may still be partially effective at allowing species to mitigate some consequences of climate change. Understanding how ongoing redistributions can allow species to retain their climatic niches is thus key for conservation^4,9,25^.

Here, we use birds as a natural system and leverage a vast trove of heterogenous citizen science data from North America to assess the patterns and drivers of climatic niche retention achieved through their geographical redistributions. We simultaneously estimate dynamics in both niche and geographical space for hundreds of bird species across multiple seasons while accounting for human sampling bias. Birds stand out as a highly mobile and intensely monitored species group with extensively documented geographical redistributions^2,3,17,18,26^. They are also central to multiple aspects of ecosystem functioning and many species are threatened by the combination of land-use and climate change^27,28^. Their mobility allows them to react to changing environments quickly and they are thus expected to be among the most successful taxa in limiting their exposure to the extensive warming the North American continent has experienced over the past 20 years.

### Realized and mitigated climatic niche change

Estimating trends and drivers of geographical redistributions and climatic niche retentions requires data at sufficiently fine spatial and temporal scales yet over large geographic extents^29^. The vast trove of observational avian citizen science data offers a unique resource that has been difficult to apply to robust trend assessments due to inherent biases resulting from where and when participants choose to record observations. To address this, we developed a novel framework that quantifies changes in species’ occurrences in space and environments conjointly with that of observer activity (Fig. 1a). Over a twenty-year period, we quantified geographical redistributions in terms of latitudinal, elevational, and distance-based movement, and climatic niche retention based on maximum daily temperatures (hereafter *MaxDailyTemp*), productivity, and precipitation levels in the 30 days leading up to each of tens of millions of records. We applied this approach to estimate niche and geographical dynamics across both the summer and winter seasons. The approach controls for any changes in observer activity that could bias the assessed spatial or climatic species metrics.

**Figure 1.**
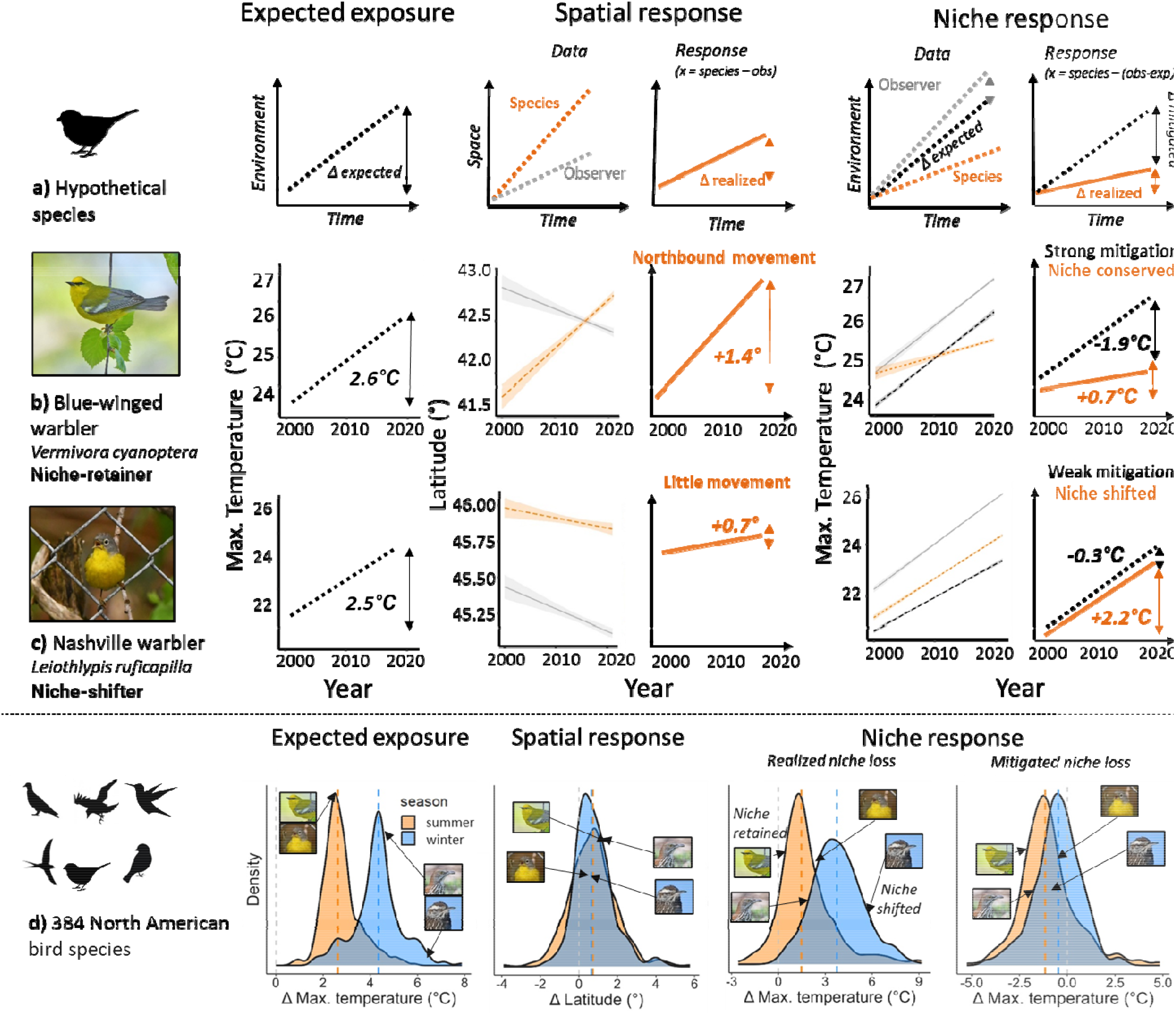
Geographic redistributions and niche loss over 20 years of climate change. Geographic redistributions and niche shifts are assessed for while controlling for trends in observer locations. The baseline trend in an environmental feature for locations where the species was observed from 2000-2004 is estimated in ‘Expected Exposure’ (black lines). In ‘Spatial Response’, trends in the location of species observations (orange) are corrected by observer trends (gray). Finally, in ‘Niche Response’, species trends in the environmental feature are corrected by baseline and observer trends, resulting in the realized niche shift for the species and mitigation of niche loss (expected – realized shift). In (a), the framework is presented for a hypothetical species. In (b-c), the framework is supported with data and estimated trends for Blue-winged warbler (*Vermivora cyanoptera*) and Nashville warbler (*Leiothlypis ruficapilla*). (d) Density plots give distributions of expected niche shift, latitudinal change, and realized and mitigated niche shift across all 384 assessed bird species in summer (orange) and winter (blue). Colored dotted lines represent corresponding seasonal means, the gray dotted line demarks zero. Example species from earlier panels and Fig. S1 are highlighted.

Consider the Blue-winged Warbler (*Vermivora cyanoptera*; Fig. 1b). In the early 2000s, it experienced an average *MaxDailyTemp* of *ca*. 23.5°C across all locations where it was observed. But over the subsequent twenty years, *MaxDailyTemp* at these same locations increased to 26.1°C, resulting in an *expected* (observer-adjusted) exposure relative to the 2000s of 2.6°C. Over the same period, the species redistributed northward by an observer-adjusted mean of 1.4° latitude (*ca*. 155 km). In estimating this value, our framework accounts for a slight southward shift in observer activity that occurred alongside the species redistribution in that region. In climatic niche space, we find that these redistributions allowed the warbler to largely compensate for substantial regional increases in temperature and by 2020 the species had a *realized MaxDailyTemp* only slightly warmer (0.7°C) than in the early 2000s, while mitigating 1.9°C of its expected exposure had it stayed put. In contrast, the Nashville Warbler (*Leiothlypis ruficapilla*; Fig. 1c) experienced a similar temperature increase in its range but did not redistribute; thus, it realized a substantial 2.2°C increase in *MaxDailyTemp* and mitigated only a minor portion of its expected exposure (0.3°C). Therefore, we consider the Blue-winged Warbler to be a ‘niche retainer’ and the Nashville Warbler a ‘niche shifter’.

Repeating this assessment for the summer distributions and climatic niches of 384 species, we find that species ranges in the year 2000 were exposed to an average 2.4 °C (+/-0.1° SE) temperature increase in the subsequent 20 years. During this period, they redistributed on average 0.73° latitude (+/-0.06°) northward, allowing them to mitigate on average 1.16 °C (+/-0.07°) of change in *MaxDailyTemp* – 44% of the expected change in exposure. However, this mitigation offered only insufficient compensation, and species still saw significant shifts in their realized *MaxDailyTemp* niches, by on average 1.47 °C (+/-0.09°; Table S1). However, these average differences hide tremendous among-species variation (Fig. 1d; Appendix 1). The majority (66%) of species experienced large realized niche shifts of >1°C, although 56% also achieved a substantial mitigation (>1°C). Only 19% of species managed to limit their summertime niche shifts to <0.5°C.

### Seasonal differences

Despite the importance of seasonality in delineating range and climatic niche boundaries in mobile organisms^30–32^, seasonality is often neglected when assessing geographical redistributions and climate change exposure^4,33,34^, and most predictions are based on once-annual surveys^3,15^. In North America, species are most likely to experience climatic niche loss during winter, which is warming faster than summer^35^. Fewer spatial constraints in individual home ranges outside the breeding season also mean potentially greater flexibility in winter to shift distributions^3,34^. However, vagile species may face greater pressure to limit exposure during the summer, when they live closest to their upper thermal limits^38^. Exploring seasonal variation can thus detect periods during which wildlife is at greatest risk of exposure^25^.

During winter, species again exhibited a range of niche-retainer and niche-shifter strategies, although given the rapid pace of winter warming, it was more difficult for species to fully retain their climatic niches and examples of niche-shifting were typically more extreme than in summer. For example, between 2000 and 2020 the Brown Thrasher (*Toxostoma rufum*) moved its wintering range north by 1.6° latitude but was only able to mitigate 2.2°C of 4.5°C expected increases in *MaxDailyTemp*, realizing a 2.3°C shift (Fig. S1a). Meanwhile, the Cactus Wren (*Campylorhynchus brunneicapillus*) only moved slightly north (0.4°) and thus realized warming of 5.3°C in *MaxDailyTemp* in only 20 years (Fig. S1b).

Across 258 species with winter ranges in the US and Canada, we find that species were exposed to 4.3 °C (+/-0.2°) increases in *MaxDailyTemp*, a much greater exposure than during summer (Fig. 1d). In response, they moved northward 0.65° (+/-0.06°) and realized 3.74 °C (+/-0.1°) increases in *MaxDailyTemp*, mitigating only 0.47 °C of warming (+/-0.07°). In winter, climatic niche loss was inevitable, as nearly all species (96%) realized niche shifts >1°C, even though 71% were able to mitigate >1°C. Therefore, birds mitigated only 11% of their expected winter exposure using movement (in contrast with 44% during summer), and only 2% of species limited their niche shift to <0.5°C. This finding supports the hypothesis that species are under greater pressures to limit their exposure to warming during the summer months, as they are living closer to their upper thermal boundaries^38^.

### Mitigating climate change exposure

We next explored the drivers of species’ realized niche shifts. We expected that while species would generally be sensitive to the degree of climate change occurring in their ranges^4,15^, species-level variability in the ability to mitigate this exposure would be best explained by recent geographical redistributions^17,18^ which may afford species an opportunity to escape exposure to climate change. However, despite the widespread assumption that species are relocating to escape the impacts of climate change and retain climatic niches ^2,3,17,18,26^, few studies have measured whether species have successfully retained their niches^19,20^.

Indeed, we find that the level of climate change exposure since 2000 strongly predicted the magnitude of niche shift subsequently realized (summer: R^2^ =0.33; p < 0.001; β = 1.3; Fig. 2a; winter: R^2^ =0.45; p < 0.001; β = 1.1; Fig. 2b). Mitigation success, measured as the difference between expected and realized exposure, was however unrelated to the original level of exposure (summer: R^2^ = 0.02; p = 1; β = 0.29; Fig. 2c; winter: R^2^ = 0.003; p = 1; β = 0.07; Fig. 2d). Thus, a species is on average expected to experience a stronger niche shift if there is more extreme climate change in their range, though with significant species-level variability.

**Figure 2.**
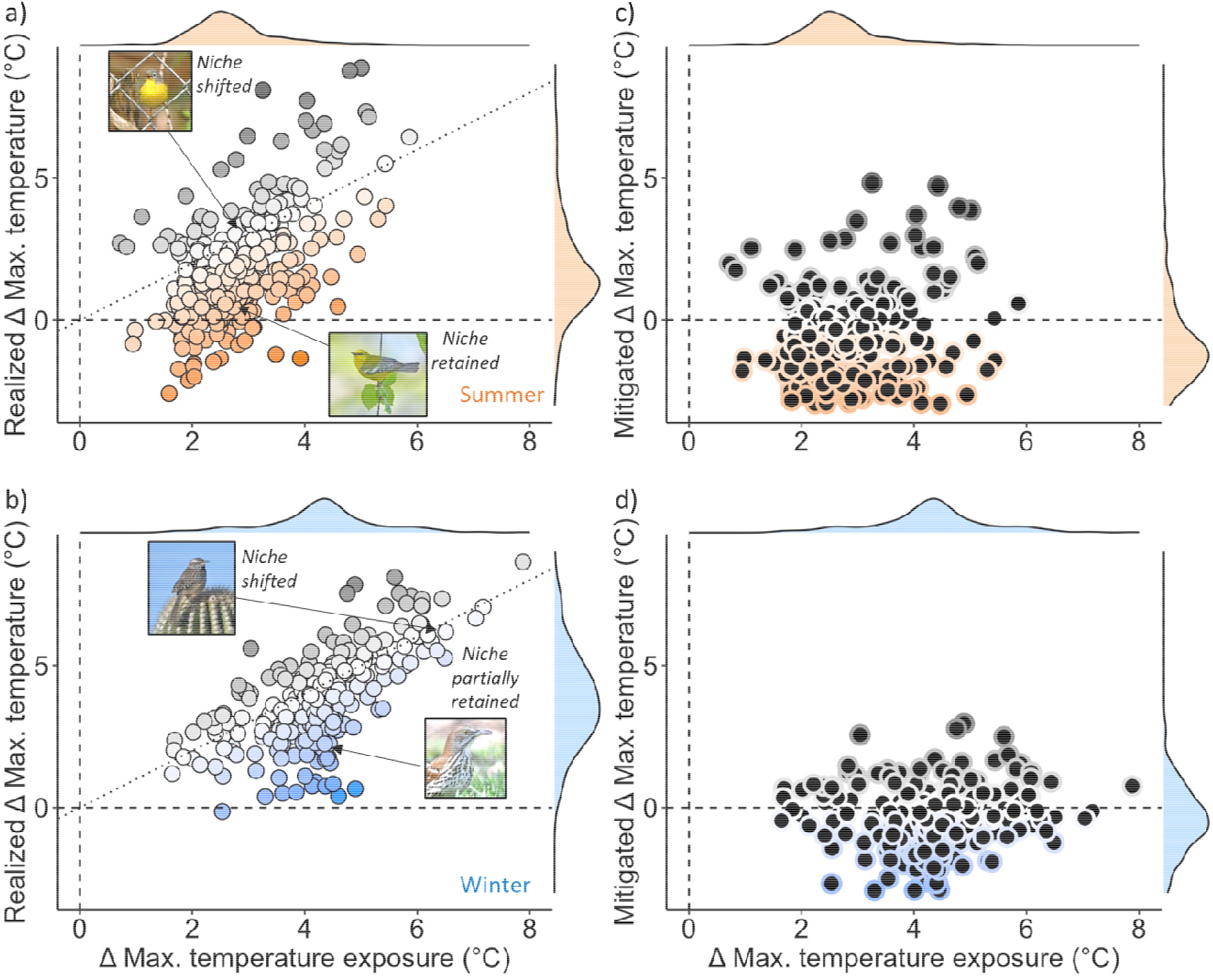
20 years of climate change exposure and niche loss across 384 North American bird species. Expected shifts in maximum daily temperature (x-axes) are compared with (a-b) realized and (c-d) mitigated shifts during (a,c) summer (orange) and (b,d) winter (blue). In (a-b), species below the dotted 1:1 line have mitigated their niche loss relative to those in their previous ranges, with darker border colors corresponding to greater distance from the line. Species on or above the line have not limited their niche loss, with darker circles representing greater exposure to temperature increases than expected based on baseline locations. The degree of mitigation is represented in (c,d; y-axes). Marginal density plots are also presented. Species covered in Figs. 1 and S1 are highlighted.

Instead, we find that individual species’ geographical redistribution (movement) is strongly associated with their respective mitigation of climate change exposure. Species that moved the furthest north experienced smaller realized increases in *MaxDailyTemp* (β = −0.26, p < 0.001; Table S2) and mitigated their exposure more than stationary species (β = −0.40, p < 0.001; Fig. 3). Movement also resulted in species occurring in less productive areas (lower EVI; β = −13.7, p < 0.001) and in areas with greater levels of precipitation (β = 0.001, p < 0.001; Fig. S2).

**Figure 3.**
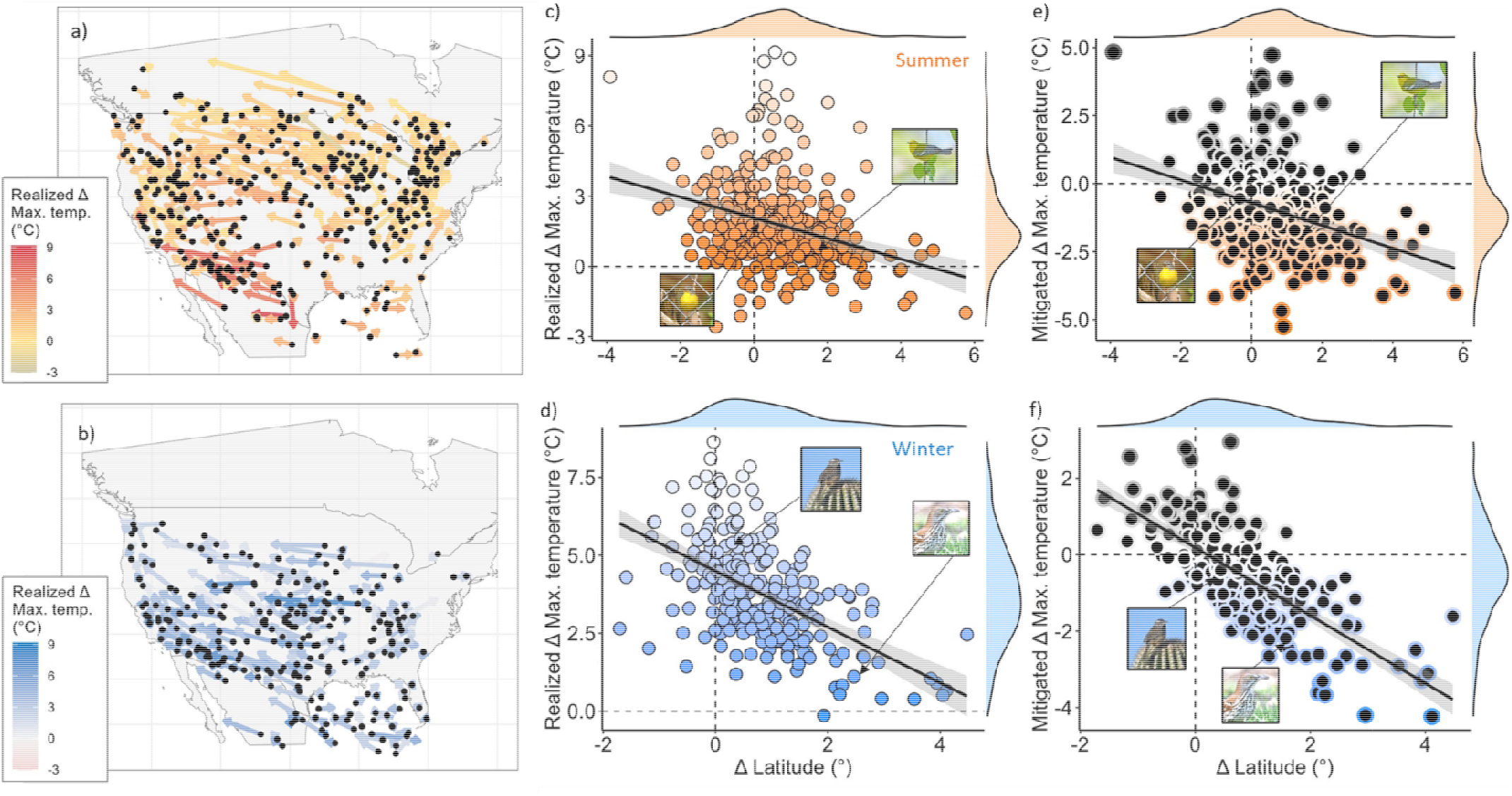
Geographic redistributions mitigated some climate change exposure. (a-b) Maps display redistributions and realized climate change exposure since 2000 for all assessed species. Black points are original (2000-2004) range centroids and arrow endpoints are recent (2016-2020) centroids based on predicted trends. Stronger colors correspond with greater realized increases in maximum temperature. (c-d) Across species, realized increases in daily max. temperature over 20 years (x-axes) are compared with latitudinal shifts (y-axes). Point color intensifies with increasing niche retention. Linear best-fit trendlines and 95% confidence intervals are shown. (e-f) Across species, mitigated increases in daily max. temperature over 20 years (x-axes) are compared with latitudinal shifts (y-axes). Border color intensifies with increasing mitigation of exposure (more negative values). All trends are visualized for (a) summer (orange) and (b) winter (blue). Dotted lines represent zero. Marginal density plots are also presented. Species covered in Figs. 1 and S1 are highlighted.

**Figure 4.**
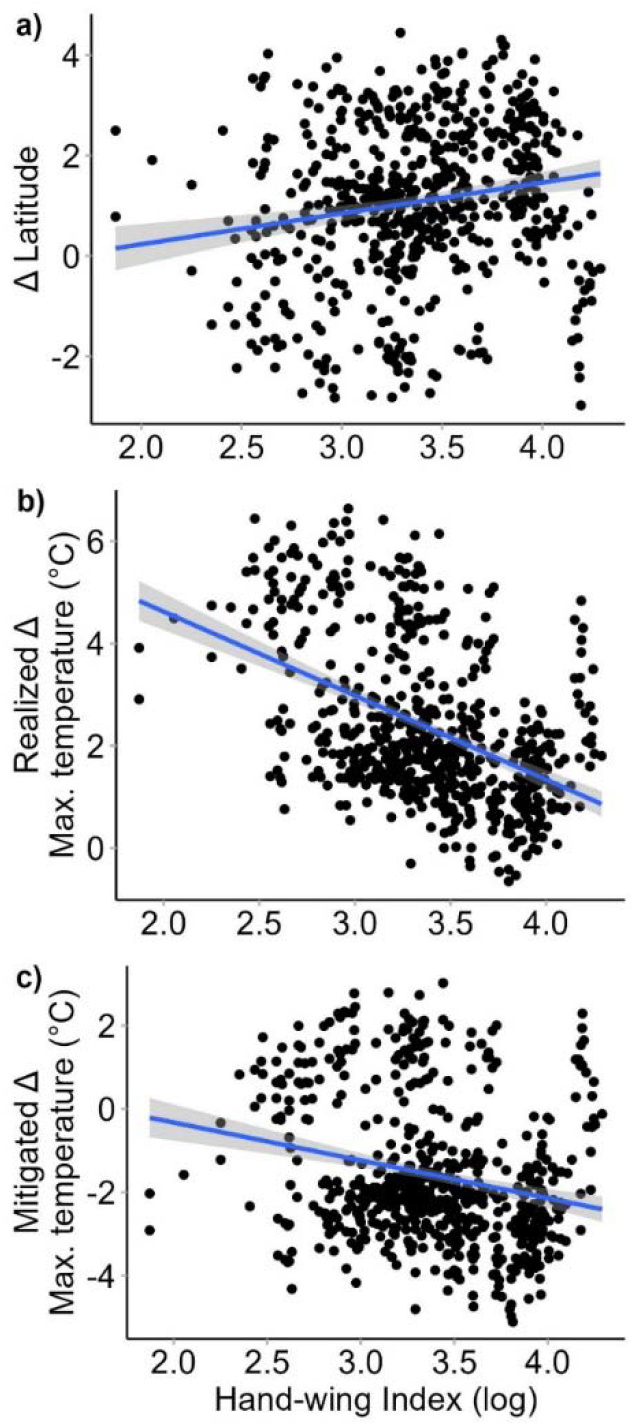
Dispersal ability is associated with range redistributions and niche loss. Partial dependence plots based on phylogenetic least-squares models visualize relationships between hand-wing index, a measure of dispersal ability, and 20-year (a) shifts in latitude, (b) realized shift in maximum temperature, (c) mitigated shift in max. temperature, while controlling for season, other functional traits, and phylogenetic structure. Points are partial residuals and lines represent the best fit through them, with gray shading indicating the confidence intervals.

Meanwhile, movement upslope was surprisingly associated with higher temperatures, although with a relatively shallow relationship (β = 12.5, p < 0.001), while distance was not (β = −1.5, p = 0.5; Figs. S3-4). Species that moved further north experienced greater reductions in productivity (β = −12.8, p < 0.001; Fig. S5) and increases in precipitation (β = 0.002, p < 0.001; Fig. S5) than expected.

Our results reveal that even though species’ geographical redistributions during the past 20 years were considerable, they only partially succeeded in allowing birds to retain their climatic niches. We see several potential causes. First, landscape fragmentation moderates the ability of species to move, and might be more consequential in certain regions (e.g., the west)^39^. The current network of protected areas does not currently provide enough connectivity to enable long-distance dispersal and avoid climate change exposure^40^; for example, Henslow’s Sparrows (*Ammodramus henslowii*) were unable to move north due to high prairie fragmentation in the central US^41^. Second, the habitat characteristics that birds depend on may not be shifting as quickly as the climate^42^, leaving some species stationary to satisfy unique habitat requirements. Third, although many bird species disperse well, some residential species (e.g., grouse) cannot disperse quickly enough to follow changing environmental conditions^43^, while migrants with high breeding site fidelity may be inherently reluctant to fly further north each year^15^.

While highly mobile species may eventually move far enough to compensate for their current climatic niche loss, movement is unlikely to keep pace with rapidly accelerating climate change^19^. Our finding that even highly mobile birds fail to keep pace with rapid climatic shifts are a cause of concern for most other taxa which are much less vagile. Species possess other tools to limit their exposure while remaining stationary. Organisms can dynamically moderate their thermal sensitivity via phenotypic plasticity^13,14^, and those with particularly fast generation times may, up to a point, be able to adapt to conditions as they change^12^. Shifts in phenological activity, including the timing of breeding or movement, towards earlier in the spring has the potential to offset climate change exposure without geographical redistributions^44,45^. Finally, microhabitats can offer animals opportunities to avoid extended exposure^46^, though with a possible cost to their ability to find food or breed^47^, and although widely recognized in its direct fitness relevance^48,49^, maximum temperatures might not always be the most relevant climatic constraint^20,50^. We highlight the importance of work that examines how species are balancing climatic niche loss across multiple environmental dimensions and across spatial scales.

### Drivers of redistributions and climatic niche retention

We next explored functional and phylogenetic drivers of geographical redistributions and climatic niche loss to uncover general relationships to that can help identify particularly vulnerable species. We expected the following species to be best at mitigating their exposure: 1) short-distance migrants and residents, given their reduced breeding and wintering site fidelity^15^; 2) species with high dispersal ability, which may easily track rapidly changing conditions in space; 3) smaller-bodied birds, which have lower thermal inertia than large-bodied birds^51,52^; and 4) habitat generalists, which may have more available habitat to move to than specialists.

Phylogeny-controlled models including all functional traits revealed that indeed, migration distance (β = −0.59, p < 0.001), body mass (β = −0.58, p < 0.001), and hand-wing index (a proxy of dispersal ability; β = 2.13, p < 0.001; Fig. 5) were associated with limiting climate change exposure via movement, although we did not detect an association with habitat generalism (Fig. S6, Tables S3-5). While species with high dispersal ability moved to mitigate temperature increases as we predicted, we found that large-bodied birds and long-distance migrants were effective at mitigating their exposure, counter to our expectations. Our findings were generally consistent when functional traits were modeled within a univariate framework (Table S6). We generally found only moderate phylogenetic signal in geographical redistributions and climatic niche loss (λ always < 0.3; k typically < 0.1; Table S7).

## Conclusions

Recent climate change is exposing wildlife to novel conditions^10,11^, causing substantial climatic niche erosion for many species^8,50^. We developed a novel approach to leverage citizen data for measuring species success in climatic niche retention. While many North American bird species have moved north, most are experiencing ever-increasing climatic niche erosion. Species that moved the furthest north, and which have the physiologically capacity to move longer distances, were most effective at limiting their exposure to climate change. However, few species were able to completely keep pace with thermal conditions in summer, and almost none could do so in winter. As the pace of climate change accelerates, the gap between the environmental conditions which species are experiencing and the conditions to which they have historically adapted will likely continue to widen, especially for species that are less mobile than birds. This finding raises important new questions about the expected consequences as this gap widens over the coming years and restructures ecological communities based on differential species movement to track climatic niches^50^. Understanding the extent, causes and consequences of geographical redistributions and climatic niche erosion will be critical for forecasting how species will fare under climate change. Our approach expands and demonstrates the role of citizen science data in supporting such a conjoint assessment of spatial and niche dynamics of biodiversity in a rapidly changing world.

## Methods

### Species selection and data

In April 2022, we accessed the Spatiotemporal Observation Annotation Tool (STOAT) v1.0, a novel cloud-based toolbox for flexible biodiversity annotations^53^, to download annotated Global Biodiversity Information Facility data (https://www.gbif.org/) between 2000-2020 for 672 bird species that annually breed or overwinter in the United States and Canada (data compilation, analyses and visualizations were all completed in R 4.1.0^54^). Our species list was based on American Birding Association birding codes^55^ updated to the Clements bird taxonomy as of 2021^56^, a widely-accepted authority to distinguish regularly occurring species from vagrants. We excluded marine species, for which weather data is unavailable, and those not native to the US or Canada.

### Observational and environmental data

Our goal was to characterize the extent to which species experienced geographical redistributions and both realized and expected niche loss over time while accounting for human observer trends. To this end, we compiled three datasets for each species: 1) species dataset, simply the observations of a given species; 2) baseline dataset, a set of annual (2000-2020) pseudo-observations corresponding to each location where the species was observed between 2000-2004, used to estimate the environmental conditions a species would have experienced had it remained stationary over the course of the study period; and 3) observer dataset, or the set of unique locations where observations of any species occurred within the 200km-buffered expert range boundary of the focal species (via https://ebird.org/science/status-and-trends/download-data, accessed April 2022), to adjust for spatial trends in observer reporting patterns.

Independent of long-term trends, spatial and temporal sampling bias are both of significant concern for point-based occurrence data, with certain places and dates oversampled relative to others, potentially biasing estimates of the species range or niches. To minimize spatiotemporal sampling bias, we thinned all three datasets by selecting one point from each 5km^2^ grid cell per week.

Though our species list was restricted to the US and Canada, we compiled data from the entirety of the American continents for each species (longitude < −25°) to be able to estimate trends across each species’ range in the entire region. Further, we temporally cropped data to season (December-February for winter and June-August for summer). We excluded species with < 20 occurrence points for either summer or winter in any single year between 2000-2020 (minimum 500 points per species across all years). To improve computational efficiency, we limited each of the three species datasets to 50,000 points apiece maximum (for species with high quantities of data) by randomly sampling points. Our final species total was 384 in summer and 258 in winter, as many species that overwinter in the tropics had insufficient data.

Using Google Earth Engine^57^, we annotated each dataset with three environmental dimensions: daily maximum temperature (MOD11A1 v006, sourced from NASA-MODIS^58^), enhanced vegetation index (EVI; from MODIS), and precipitation (from CHELSA 2.1^59^), each summarized to a 1km buffer (0.5km radius) over 30 days prior to the observation (imported using *jsonlite* and *httr* packages^60,61^). We characterized the location of each point with latitude, elevation, and distance, which is the absolute value of great circle distance from the 2000-2004 centroid for the species (i.e., distance from original range center), calculated using the *fields* package^62^.

Elevation was annotated from the Global Multi-Terrain Elevation Dataset, a product of USGS and the National Geospatial-Intelligence Agency^63^ using the *raster* package^64^. All data management was completed using *tidyverse* packages^65^. For each species, we restricted all observations or pseudo-observations to those with environmental data available for all dimensions, which eliminated only those in some coastal areas and small islands (<0.3% of the data).

### Statistical framework

Our first goal was to understand how species redistributed their ranges over 20 years of climate change. We characterized observation-level changes in the distributions of three spatial metrics: latitude, elevation, and distance. For every species, we first combined the species and observer datasets then fit generalized linear models (GLMs) to calculate the coefficient of the interaction between year and data type, a two-level categorical variable describing whether the point is ‘species’ or ‘observer’ (as the contrast), with a spatial metric as the response variable. Species and observer datasets were equalized in size to ensure equal representation in the model by randomly subsampling points from the smaller dataset.

We fit three models per species, one with each metric as a response variable. Thus, the interaction coefficient reflects the difference between the ‘species’ (β_sp_) and ‘observer’ (β_obs_) slopes of a spatial metric. A high value suggests that the species moved north, upslope, or further from its original location relative to observers in the region, while a neutral value suggests little movement. Hereafter, we refer to these interaction coefficients as ‘geographical redistributions’. When considering elevation, we limited species to those mainly found in the western US and Canada (>75% of range in < −100° longitude), for which elevation is most relevant (summer n = 96, winter n = 57).

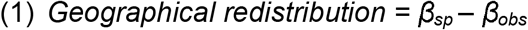

Our second goal was to quantify the extent of niche shift over this period. We quantified three measures of niche loss for each species and season: 1) expected niche shift, or the amount of change in the niche dimension in areas where the species was observed from 2000-2004; 2) realized (actual) niche shift, or the change in the niche dimension in areas where the species was observed; and 3) mitigated niche shift, or the difference between the expected and realized niche shift, representing the amount of niche space the species avoided losing via movement. For every species, we merged the species, observer, and baseline datasets (equalizing them in size as above), then fit GLMs with one niche component as the response variable (max. temperature, EVI, or precipitation) and interacting year with data type. We fit three models per species, one with each niche component as a response variable.

Using the *emmeans* package^66^, we compiled the coefficients for the species, observer, and baseline datasets from the model, reflecting the temporal shift for each. The slope coefficient among baseline points (β_base_) is the *expected niche shift*, reflecting climate change in the region. Our goal was to adjust for shifts in niches driven solely by observer movement, but not the climate change baked into the observer trends; thus, we calculated a correction factor equaling the difference between observer (β_obs_) and baseline (i.e., climate change) coefficients. Thus, we removed the climate change component from the change in niche dimension associated with observer movement, and the correction factor represents the amount of niche space driven by observer movement over the 20 years. We then calculated the difference between β_sp_ and the correction factor to find the *realized niche loss*. A greater value suggests that the species experienced increasing temperatures, EVI, or precipitation, a neutral value suggests little niche change, and a negative value suggests a decrease.

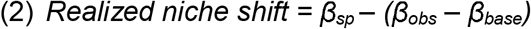

Our third goal was to understand how species mitigated their expected niche loss. To calculate *mitigated niche loss*, we subtracted expected niche loss from realized niche loss. A negative value suggests that the species experienced reduced temperatures, EVI, or precipitation relative to expected, a neutral value suggests that the species experienced similar climate change, and a positive value suggests an increase.

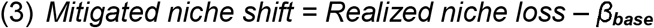

To summarize, for every species and season we compiled twelve values: geographical redistributions (latitude, elevation, or distance) and expected, realized, or mitigated niche shift (in temperature, EVI, or precipitation). To evaluate whether the magnitude and direction of the geographical redistribution is associated with niche shift or mitigated niche shift at the species level, we fit generalized linear models for every pairwise combination (18 models). We created all data visualizations using *ggplot2*^67^.

### Functional species traits and phylogeny

We obtained species-level functional trait values and an avian phylogenetic tree to test functional and phylogenetic associations with geographical redistributions and niche loss. We derived migration distances from^68; La Sorte personal communication^, and body mass and hand-wing index from AVONET^69^. Further, we calculated species-level landcover diversity index (following^31^) to represent habitat generalism, based on mean partial effects of all landcover covariates in independent continent-wide species distribution models (Cohen and Jetz *in review*). We updated an avian phylogeny from Jetz et al.^70^ to account for recent taxonomic changes, updating species names to the Clements as of 2021^71^ and treating more recently split species as having no phylogenetic distance (Table S1). We also harmonized all species names to Clements.

We fit nine multivariate phylogenetic generalized least-squares (PGLS) models at the species level to assess the simultaneous influence of traits and phylogeny on each measure of geographical redistribution, realized niche loss, or mitigated niche loss. Phylogenetically correlated model errors in PGLS account for the non-independence of species due to their phylogenetic relatedness^72^. The dependence of model errors arises from trait axes that we did not include in the analysis and that may be subject to niche conservatism so that model errors reflect the unobserved traits and thus phylogenetic distance between species. We included the four functional traits as predictor variables (log-transforming body mass and hand-wing index) and controlled for phylogenetic structure. We generated partial residual plots relating each trait to range or niche shifts based on model predictions.

To assess phylogenetic correlation among geographical redistributions and niche shifts, we calculated Blomberg’s K^73^ and lambda^74^ for each of the nine measures of geographical redistribution, realized niche loss, or mitigated niche loss, comparing each to null distributions after randomizing species’ responses 1,000 times using the *picante* package^75^.

## Notes

https://github.com/jeremy-cohen/changing-seasonal-niches

### Competing Interest Statement

The authors have declared no competing interest.

