## supplement for "Geographic redistributions are insufficient to mitigate the erosion of species’ environmental niches"

**Supplementary Information**

**Appendix 1.** Estimates of realized and mitigated niche loss and geographical redistributions between 2000-2020 for all species during summer and winter.

**Table S1.** Mean (+/- SE) estimated realized and mitigated niche loss and geographical redistributions across species for summer and winter.

|  |  | Summer | | Winter | |
| --- | --- | --- | --- | --- | --- |
| Variable | Trend | Mean | SE | Mean | SE |
| Max. Temperature | realized | 1.468 | 0.09 | 3.742 | 0.1 |
| Max. Temperature | mitigated | -1.161 | 0.075 | -0.465 | 0.075 |
| EVI | realized | -0.01 | 0.001 | -0.005 | 0.001 |
| EVI | mitigated | -0.018 | 0.001 | -0.009 | 0.001 |
| Precipitation | realized | 102.137 | 4.645 | 158.486 | 7.437 |
| Precipitation | mitigated | -15.347 | 3.034 | 34.484 | 5.268 |
| Latitude |  | 0.726 | 0.062 | 0.646 | 0.061 |
| Elevation |  | 39.164 | 6.019 | -5.762 | 4.218 |
| Distance |  | 66.294 | 4.946 | 50.115 | 6.296 |

**Table S2.** Results of generalized linear models comparing geographical redistributions with realized or mitigated niche loss across species. R^2^ values are included.

| Niche variable | Spatial variable | Coefficient | SD | p-value | R^2^ value |
| --- | --- | --- | --- | --- | --- |
| Realized Max. Temperature | Latitude | -0.262 | 0.025 | 0 | 0.155 |
| Realized Max. Temperature | Elevation | 12.449 | 2.355 | 0 | 0.15 |
| Realized Max. Temperature | Distance | -1.451 | 2.221 | 0.514 | 0.002 |
| Realized EVI | Latitude | -13.724 | 1.876 | 0 | 0.08 |
| Realized EVI | Elevation | -410.533 | 175.089 | 0.019 | 0.12 |
| Realized EVI | Distance | -1000.826 | 157.258 | 0 | 0.063 |
| Realized Precipitation | Latitude | 0.001 | 0 | 0.001 | 0.017 |
| Realized Precipitation | Elevation | -0.083 | 0.039 | 0.033 | 0.118 |
| Realized Precipitation | Distance | -0.022 | 0.036 | 0.55 | 0.002 |
| Mitagated Max. Temperature | Latitude | -0.396 | 0.029 | 0 | 0.226 |
| Mitagated Max. Temperature | Elevation | 15.326 | 2.942 | 0 | 0.149 |
| Mitagated Max. Temperature | Distance | 4.01 | 2.769 | 0.148 | 0.005 |
| Mitagated EVI | Latitude | -12.77 | 1.964 | 0 | 0.064 |
| Mitagated EVI | Elevation | -701.767 | 180.346 | 0 | 0.133 |
| Mitagated EVI | Distance | -1140.827 | 162.121 | 0 | 0.075 |
| Mitagated Precipitation | Latitude | 0.002 | 0.001 | 0 | 0.025 |
| Mitagated Precipitation | Elevation | -0.14 | 0.056 | 0.014 | 0.12 |
| Mitagated Precipitation | Distance | -0.107 | 0.052 | 0.041 | 0.008 |

**Table S3.** Results of phylogenetic generalized least-squares models testing associations between functional traits and niche loss across species.

| Realized Max. Temperature | Value | Std.Error | t-value | p-value |
| --- | --- | --- | --- | --- |
| (Intercept) | 5.448 | 1.571 | 3.468 | 0.001 |
| Migration.Distance | -0.468 | 0.026 | -18.229 | <0.001 |
| Body.Mass | -0.459 | 0.109 | -4.214 | <0.001 |
| LDI | -1.988 | 0.837 | -2.376 | 0.018 |
| Hand.Wing.Index | 0.733 | 0.377 | 1.944 | 0.052 |
| season (winter) | 1.007 | 1.054 | 0.955 | 0.340 |
| Realized EVI | Value | Std.Error | t-value | p-value |
| (Intercept) | 0.009 | 0.025 | 0.347 | 0.729 |
| Migration.Distance | 0.006 | 0 | 14.853 | <0.001 |
| Body.Mass | 0.006 | 0.002 | 3.694 | <0.001 |
| LDI | -0.051 | 0.013 | -3.845 | <0.001 |
| Hand.Wing.Index | -0.024 | 0.006 | -3.985 | <0.001 |
| season (winter) | 0.039 | 0.017 | 2.324 | 0.020 |
| Realized Precipitation | Value | Std.Error | t-value | p-value |
| (Intercept) | 23.688 | 94.423 | 0.251 | 0.802 |
| Migration.Distance | -0.961 | 1.542 | -0.623 | 0.533 |
| Body.Mass | 6.5 | 6.551 | 0.992 | 0.322 |
| LDI | 56.596 | 50.28 | 1.126 | 0.261 |
| Hand.Wing.Index | 13.932 | 22.678 | 0.614 | 0.539 |
| season (winter) | 59.599 | 63.362 | 0.941 | 0.347 |

**Table S4.** Results of phylogenetic generalized least-squares models testing associations between functional traits and mitigated niche loss across species.

| Max. Temperature (mitigated) | Value | Std.Error | t-value | p-value |
| --- | --- | --- | --- | --- |
| (Intercept) | -2.766 | 1.625 | -1.702 | 0.089 |
| Migration.Distance | -0.591 | 0.027 | -22.251 | <0.001 |
| Body.Mass | -0.578 | 0.113 | -5.129 | <0.001 |
| LDI | 0.29 | 0.865 | 0.335 | 0.738 |
| Hand.Wing.Index | 2.133 | 0.39 | 5.464 | <0.001 |
| season (nonbreeding) | -0.883 | 1.09 | -0.81 | 0.418 |
| EVI (mitigated) | Value | Std.Error | t-value | p-value |
| (Intercept) | 0.021 | 0.026 | 0.806 | 0.420 |
| Migration.Distance | 0.008 | 0 | 18.569 | <0.001 |
| Body.Mass | 0.01 | 0.002 | 5.431 | <0.001 |
| LDI | -0.067 | 0.014 | -4.758 | <0.001 |
| Hand.Wing.Index | -0.034 | 0.006 | -5.325 | <0.001 |
| season (nonbreeding) | 0.04 | 0.018 | 2.281 | 0.023 |
| Precipitation (mitigated) | Value | Std.Error | t-value | p-value |
| (Intercept) | 2.777 | 69.306 | 0.04 | 0.968 |
| Migration.Distance | 16.852 | 1.132 | 14.887 | <0.001 |
| Body.Mass | 29.453 | 4.809 | 6.125 | <0.001 |
| LDI | 32.983 | 36.905 | 0.894 | 0.372 |
| Hand.Wing.Index | -76.831 | 16.646 | -4.616 | <0.001 |
| season (nonbreeding) | 81.585 | 46.507 | 1.754 | 0.080 |

**Table S5.** Results of phylogenetic generalized least-squares models testing associations between functional traits and geographical redistributions across species.

| Latitude | Value | Std.Error | t-value | p-value |
| --- | --- | --- | --- | --- |
| (Intercept) | -0.312 | 1.433 | -0.218 | 0.827 |
| Migration.Distance | 0.462 | 0.023 | 19.741 | <0.001 |
| Body.Mass | 0.429 | 0.099 | 4.314 | <0.001 |
| LDI | 2.820 | 0.763 | 3.696 | <0.001 |
| Hand.Wing.Index | -1.675 | 0.344 | -4.867 | <0.001 |
| season (nonbreeding) | 1.715 | 0.961 | 1.784 | 0.075 |
| Elevation | Value | Std.Error | t-value | p-value |
| (Intercept) | 480.587 | 91.627 | 5.245 | <0.001 |
| Migration.Distance | 10.743 | 1.497 | 7.178 | <0.001 |
| Body.Mass | 4.550 | 6.357 | 0.716 | 0.474 |
| LDI | -133.287 | 48.791 | -2.732 | 0.006 |
| Hand.Wing.Index | -106.195 | 22.007 | -4.826 | <0.001 |
| season (nonbreeding) | -114.804 | 61.486 | -1.867 | 0.062 |
| Distance | Value | Std.Error | t-value | p-value |
| (Intercept) | 24.277 | 82.607 | 0.294 | 0.769 |
| Migration.Distance | -1.381 | 1.349 | -1.024 | 0.306 |
| Body.Mass | -12.227 | 5.732 | -2.133 | 0.033 |
| LDI | 97.558 | 43.988 | 2.218 | 0.027 |
| Hand.Wing.Index | 36.951 | 19.84 | 1.862 | 0.063 |
| season (nonbreeding) | -69.333 | 55.433 | -1.251 | 0.211 |

**Table S6.** Results of univariate phylogenetic generalized least-squares models testing associations between a single functional trait and niche shifts or geographical redistributions across species.

| Model | Coefficient | p-value | R^2^ value |
| --- | --- | --- | --- |
| tmax_Migration.Distance | -0.467 | 0 | 0.355 |
| tmax_Body.Mass | -0.902 | 0 | 0.266 |
| tmax_LDI | -1.171 | 0.283 | 0.267 |
| tmax_Hand.Wing.Index | -2.72 | 0 | 0.284 |
| tmax_mitigated_Migration.Distance | -0.554 | 0 | 0.082 |
| tmax_mitigated_Body.Mass | -1.109 | 0 | 0.046 |
| tmax_mitigated_LDI | 1.496 | 0.217 | 0.034 |
| tmax_mitigated_Hand.Wing.Index | -2.16 | 0 | 0.034 |
| modisevi_Migration.Distance | 0.006 | 0 | 0.022 |
| modisevi_Body.Mass | 0.012 | 0 | 0.038 |
| modisevi_LDI | -0.064 | 0 | 0.006 |
| modisevi_Hand.Wing.Index | 0.019 | 0.003 | 0.046 |
| modisevi_mitigated_Migration.Distance | 0.008 | 0 | 0.053 |
| modisevi_mitigated_Body.Mass | 0.018 | 0 | 0.069 |
| modisevi_mitigated_LDI | -0.086 | 0 | 0.023 |
| modisevi_mitigated_Hand.Wing.Index | 0.024 | 0.001 | 0.076 |
| prec_Migration.Distance | -0.24 | 0.859 | 0.055 |
| prec_Body.Mass | 5.445 | 0.388 | 0.056 |
| prec_LDI | 53.83 | 0.28 | 0.057 |
| prec_Hand.Wing.Index | 12.361 | 0.538 | 0.056 |
| prec_mitigated_Migration.Distance | 15.956 | 0 | 0.144 |
| prec_mitigated_Body.Mass | 43.179 | 0 | 0.148 |
| prec_mitigated_LDI | -14.677 | 0.743 | 0.156 |
| prec_mitigated_Hand.Wing.Index | 53.739 | 0.003 | 0.148 |
| lat_Migration.Distance | 0.431 | 0 | 0.026 |
| lat_Body.Mass | 0.803 | 0 | 0.001 |
| lat_LDI | 1.895 | 0.06 | 0.046 |
| lat_Hand.Wing.Index | 1.742 | 0 | 0.025 |
| elev_Migration.Distance | 7.79 | 0 | 0.125 |
| elev_Body.Mass | 12.28 | 0.056 | 0.114 |
| elev_LDI | -160.069 | 0.002 | 0.123 |
| elev_Hand.Wing.Index | -34.428 | 0.093 | 0.125 |
| dist_Migration.Distance | -0.911 | 0.445 | 0.012 |
| dist_Body.Mass | -13.043 | 0.019 | 0.008 |
| dist_LDI | 114.204 | 0.009 | 0.001 |
| dist_Hand.Wing.Index | 23.243 | 0.19 | 0.033 |

**Table S7.** Results of tests for phylogenetic signal (Lambda and Blomberg’s K) in geographical redistributions, niche shifts, or mitigated niche shifts across species.

| Metric | Lambda | p-value | K | p-value |
| --- | --- | --- | --- | --- |
| Max. Temperature | 0.231 | 0 | 2.00E-04 | 0.218 |
| Mitigated Max. Temperature | 0.263 | 0 | 0 | 0.065 |
| EVI | 0.266 | 0 | 0 | 0.004 |
| Mitigated EVI | 0.29 | 0 | 0 | 0.001 |
| Precipitation | 0.028 | 0.463 | 0.4627 | 0.406 |
| Mitigated Precipitation | 0.237 | 0.004 | 0.0037 | 0.003 |
| Latitude | 0.196 | 0 | 3.00E-04 | 0.031 |
| Elevation | 0 | 1 | 1 | 0.124 |
| Distance | 0 | 1 | 1 | 0.003 |

**Supplementary figures**

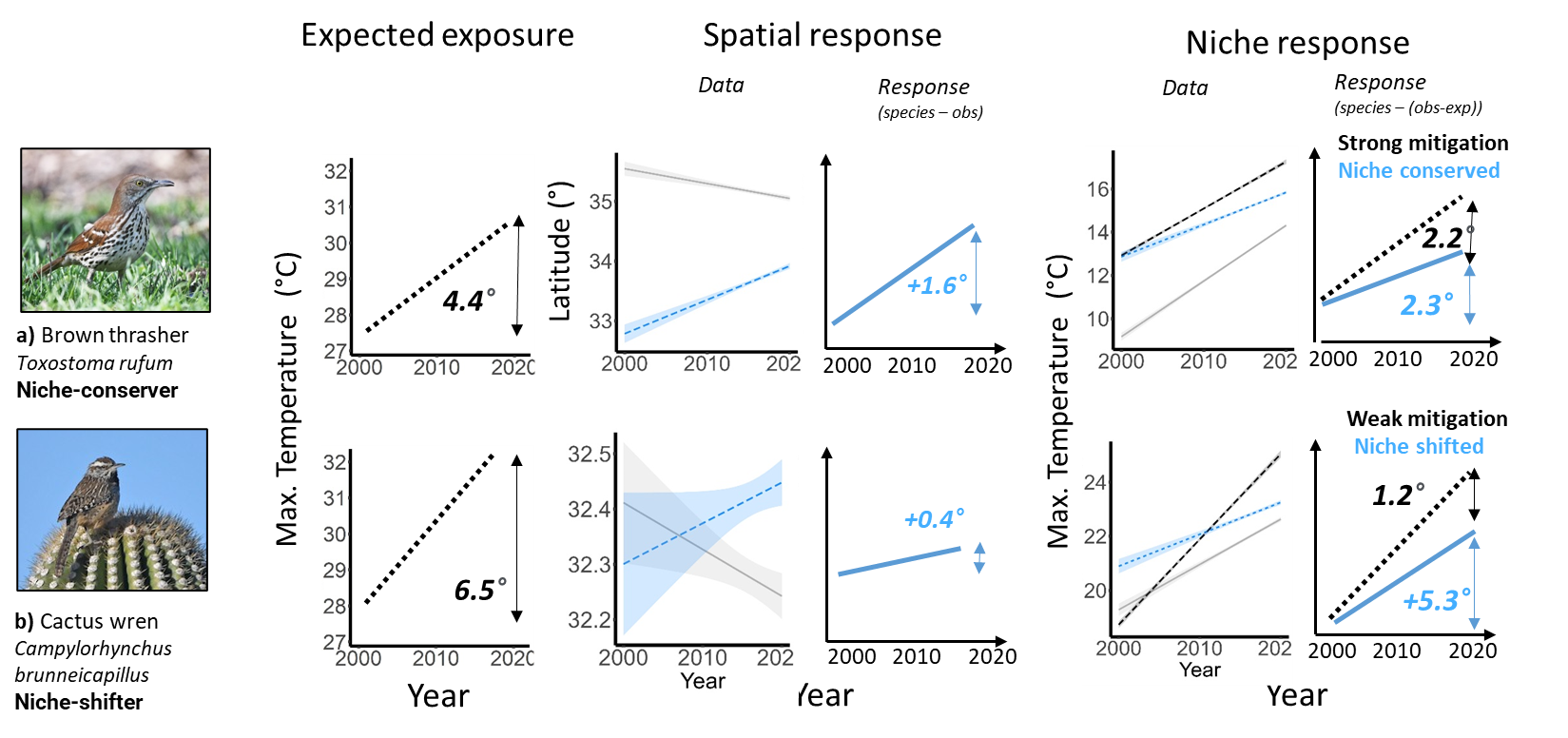

**Figure S1. Winter examples of spatial redistributions and niche loss over 20 years of climate change.** We present an approach to quantify geographical redistributions and niche loss for wildlife species using observational data. The overall trend in the environmental feature for ‘original’ locations is estimated in ‘Expected Exposure’ (black lines). In ‘Spatial Response’, trends in the location of species observations (blue) are adjusted by observer trends (gray), resulting in the spatial response. Finally, in ‘Niche Response’, trends in the environmental feature at species points are adjusted by trends at ‘original’ locations and observers, resulting in the realized niche loss for the species and mitigation of niche loss (expected – realized loss). In (a-b), the framework is supported with data and estimated trends for brown thrasher (*Toxostoma rufum*) and cactus wren (*Campylorhynchus brunneicapillus*), respectively.

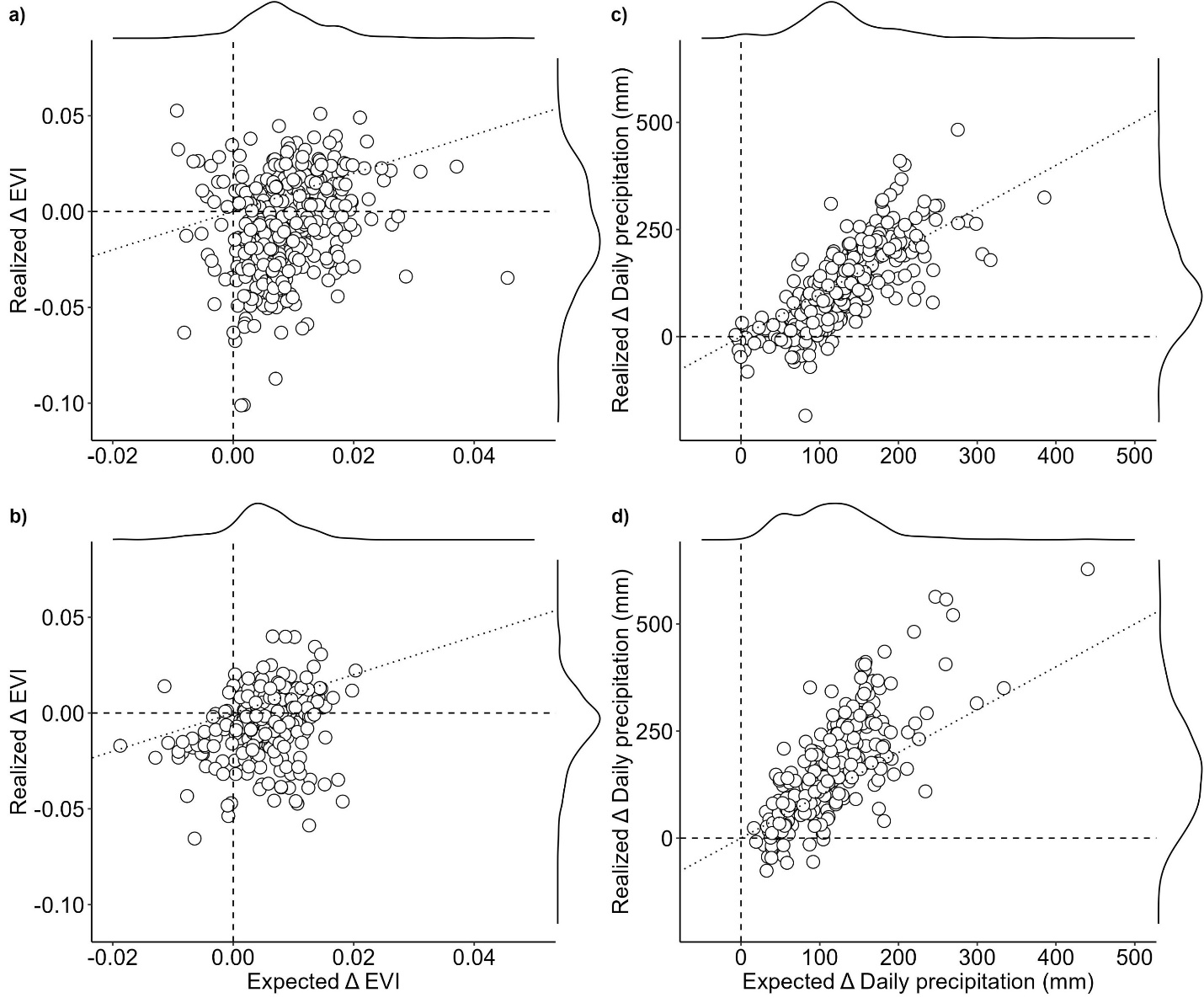
 **Figure S2. Climate change exposure and niche loss among North American birds over the past 20 years.** Across species, expected changes in (a-b) EVI or (c-d) daily precipitation (mm; x-axes) are compared with (a-b) realized changes during (a,c) summer and (b,d) winter. Species closer to zero on the y-axis have mitigated their niche loss. Species farther from zero have experienced greater niche loss.

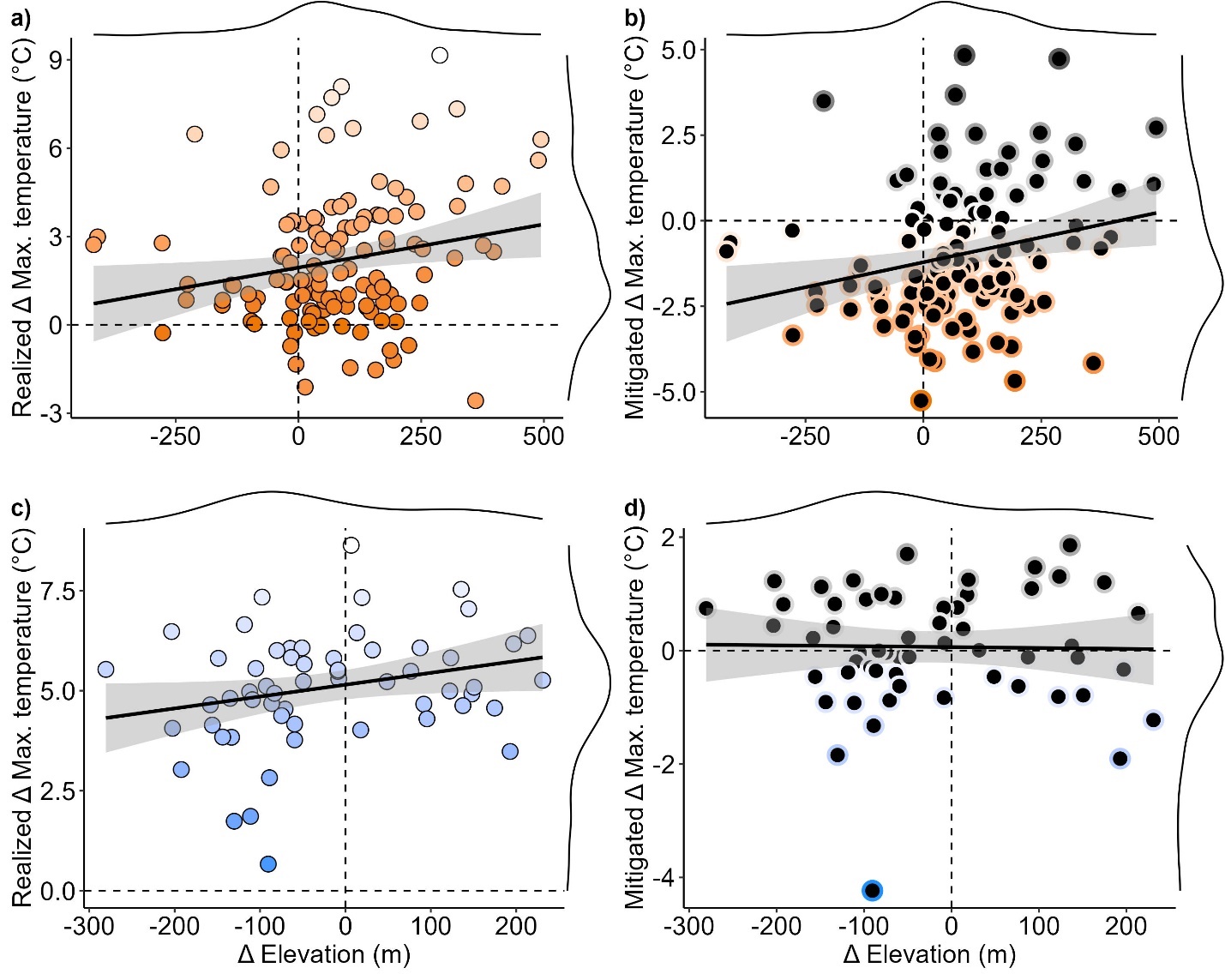

**Figure S3. Relationships between elevational shift and realized niche loss during summer and winter.** Across species, elevational shifts (x-axes) are compared with (a-b) realized or (c-d) mitigated increases in daily max. temperature over 20 years (y-axes). Linear best-fit trendlines and 95% confidence intervals are shown. Border color intensifies with increasing mitigation of exposure (more negative values). All trends are visualized for (a) summer (orange) and (b) winter (blue). Dotted lines represent zero lines.

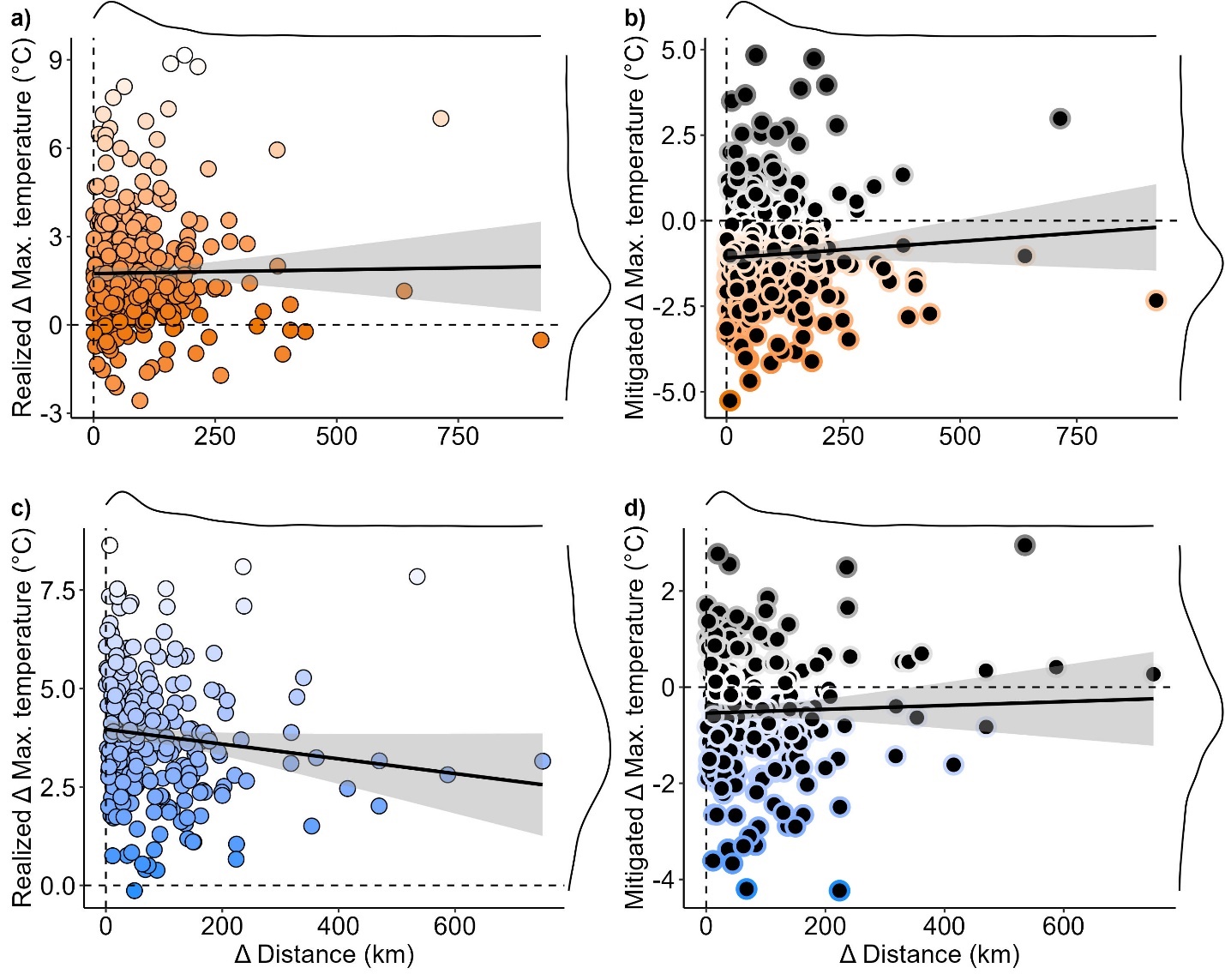

**Figure S4. Relationships between distance moved and niche loss during summer and winter.** Across species, distance moved (x-axes) is compared with (a-b) realized or (c-d) mitigated increases in daily max. temperature over 20 years (y-axes). Linear best-fit trendlines and 95% confidence intervals are shown. Border color intensifies with increasing mitigation of exposure (more negative values). All trends are visualized for (a) summer (orange) and (b) winter (blue). Dotted lines represent zero lines.

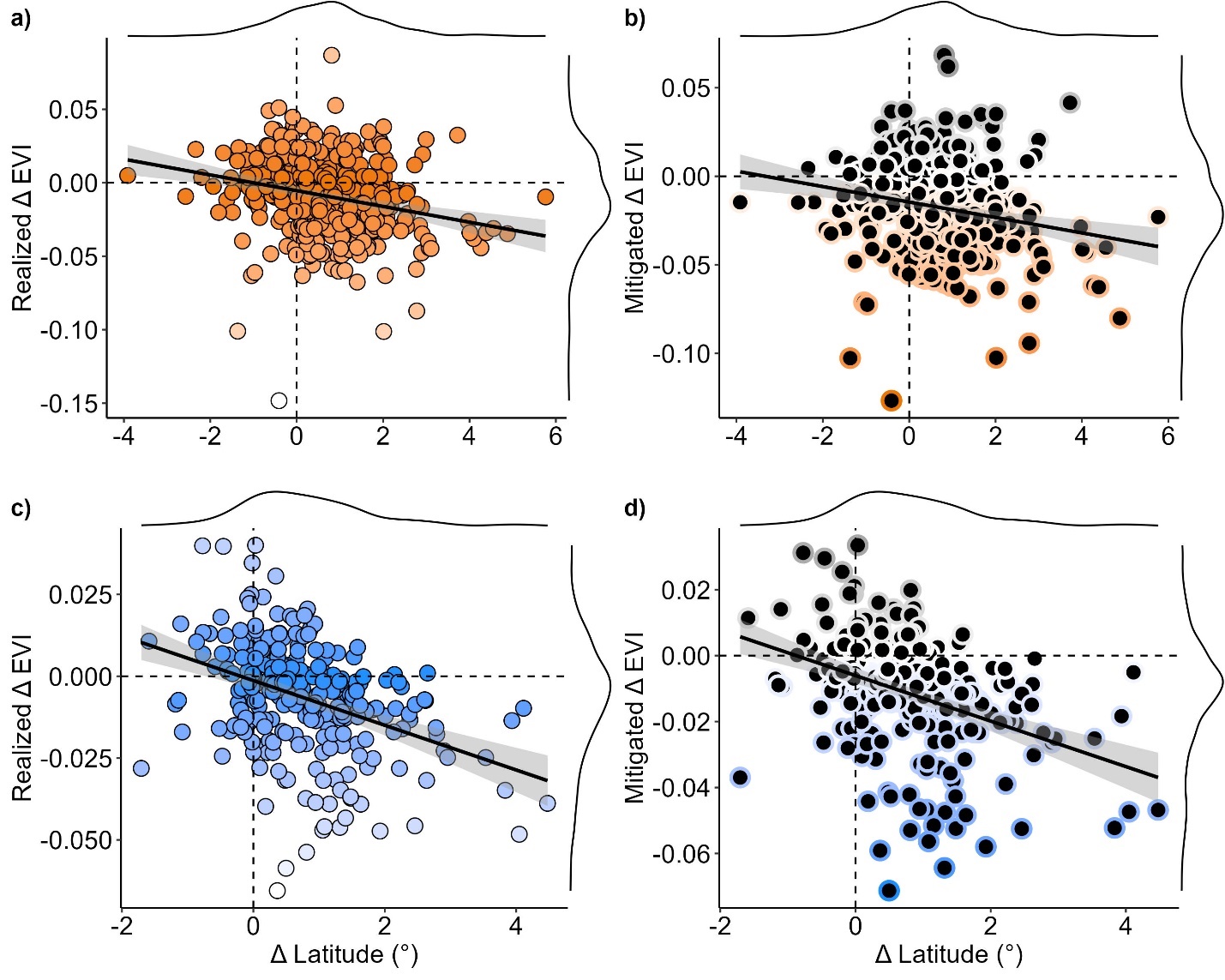

**Figure S5. Relationships between distance moved and shift in EVI during summer and winter.** Across species, distance moved (x-axes) is compared with (a-b) realized or (c-d) mitigated increases in EVI over 20 years (y-axes). Linear best-fit trendlines and 95% confidence intervals are shown. Border color intensifies with increasing mitigation of exposure (more negative values). All trends are visualized for (a) summer (orange) and (b) winter (blue). Dotted lines represent zero lines.

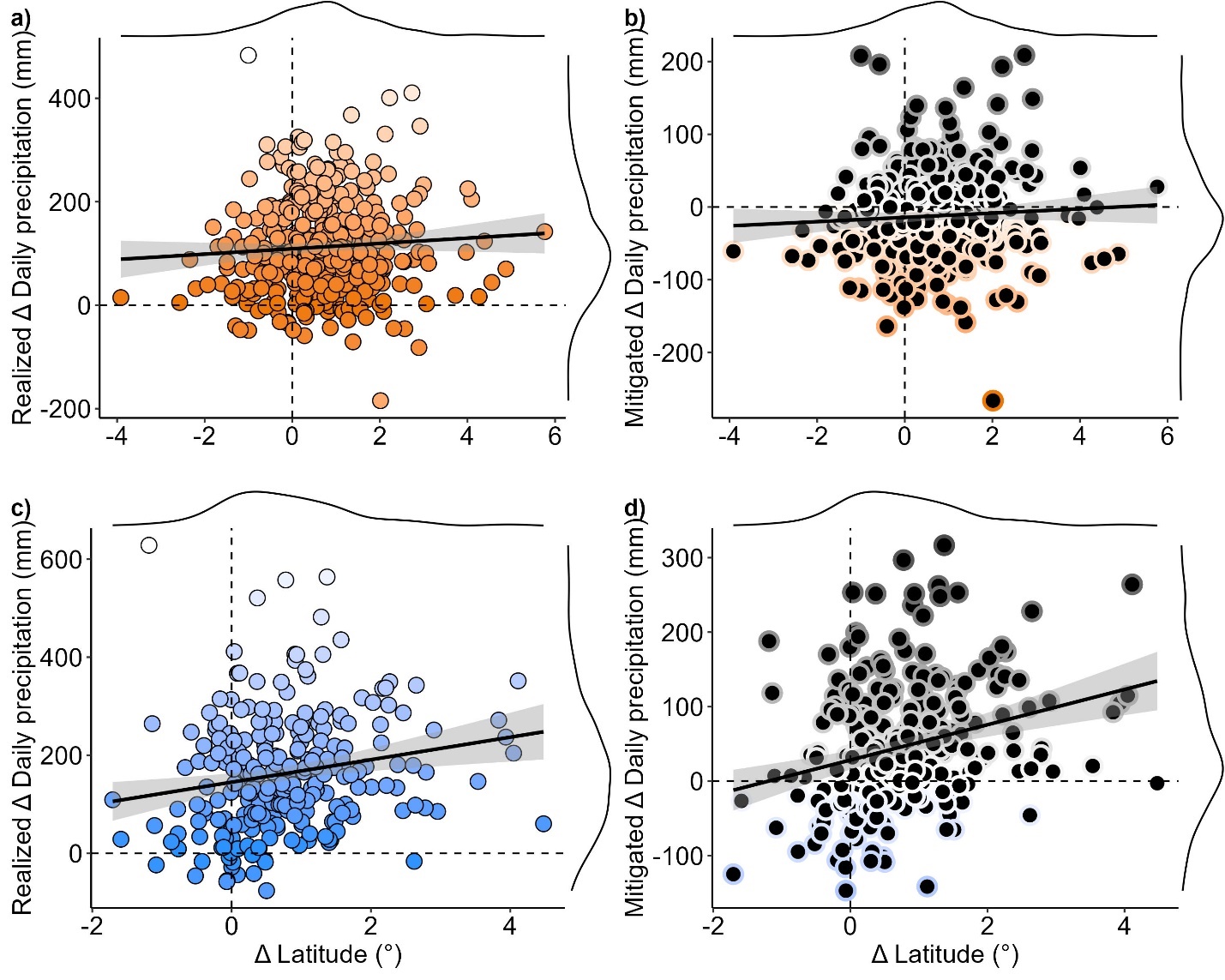

**Figure S6. Relationships between distance moved and shift in precipitation during summer and winter.** Across species, distance moved (x-axes) is compared with (a-b) realized or (c-d) mitigated increases in daily precipitation (mm) over 20 years (y-axes). Linear best-fit trendlines and 95% confidence intervals are shown. Border color intensifies with increasing mitigation of exposure (more negative values). All trends are visualized for (a) summer (orange) and (b) winter (blue). Dotted lines represent zero lines.

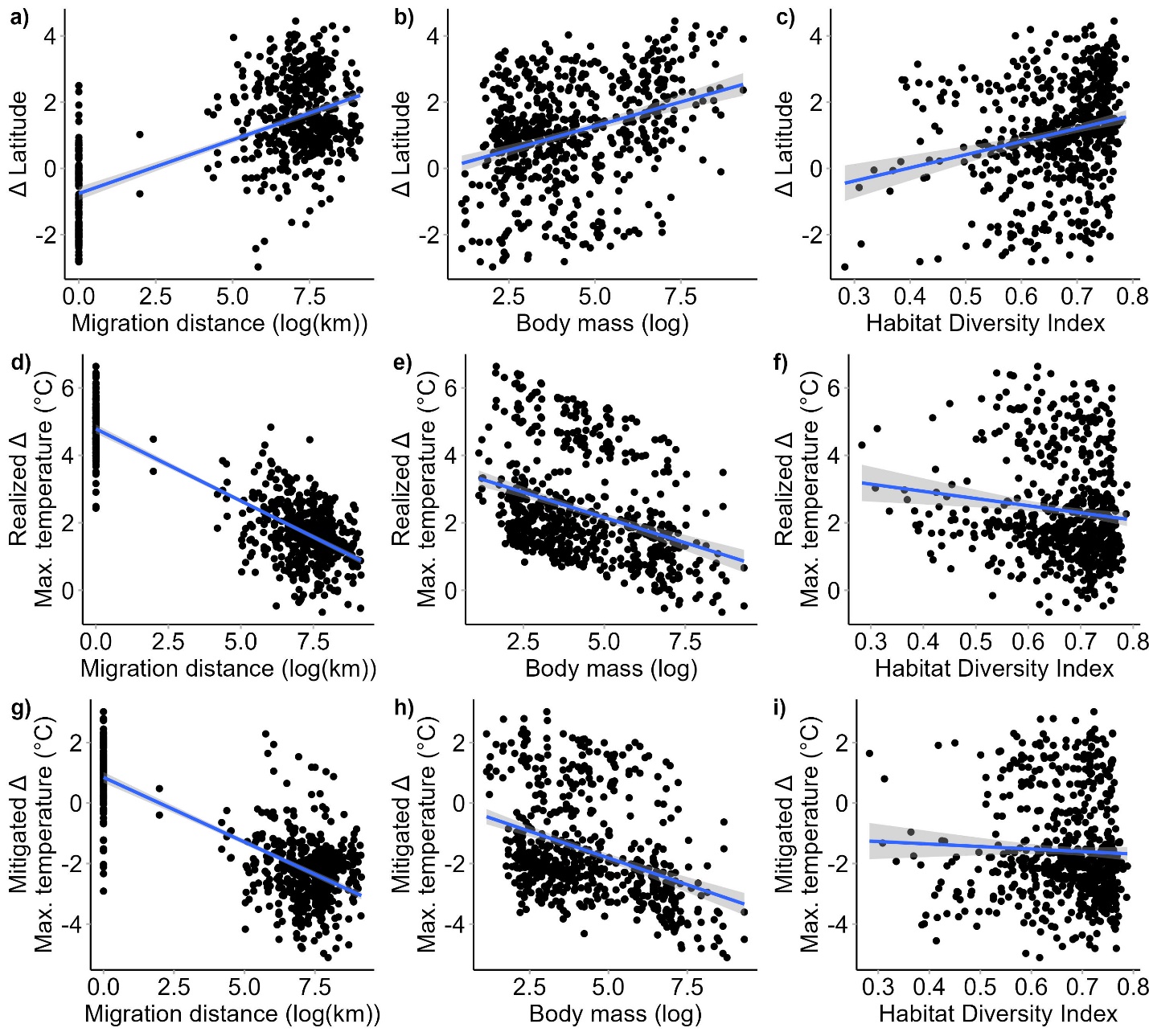
**Figure S7. Species’ functional traits are associated with geographic redistributions and niche loss.** Partial dependence plots visualize relationships between 20-year shifts in (a-c) latitude, (d-f) realized shift in max. temperature, and (g-i) mitigated shift in max. temperature, compared with species functional traits – (a,d,g) migration distance, (b,e,h) body size, and (c,f,i) habitat diversity index, a measure of habitat generalism. Trendlines are derived from phylogenetic least-squares models and represent line of best fit, with gray shading representing associated confidence intervals. Points are partial residuals for each species.
